# Still no evidence for disruption of global patterns of nest predation in shorebirds

**DOI:** 10.1101/2021.02.17.431576

**Authors:** Martin Bulla, Mihai Valcu, Bart Kempenaers

## Abstract

Many shorebird species are rapidly declining (Piersma *et al*. 2016; Munro 2017; Studds *et al*. 2017), but it is not always clear why. Deteriorating and disappearing habitat, e.g. due to intensive agriculture (Donal *et al*. 2001; Kentie *et al*. 2013; Kentie *et al*. 2018), river regulation (Nebel *et al*. 2008) or mudflat reclamation (Ma *et al*. 2014; Larson 2017), and hunting (Reed *et al*. 2018; Gallo-Cajiao *et al*. 2020) are some of the documented causes. A recent study suggests yet another possible cause of shorebird decline: a global increase in nest predation (Kubelka *et al*. 2018). The authors compiled an impressive dataset on patterns of nest predation in shorebirds and their analyses suggest that global patterns of nest predation have been disrupted by climate change, particularly in the Arctic. They go as far as to conclude that the Arctic might have become an ecological trap (Kubelka *et al*. 2018). Because these findings might have far-reaching consequences for conservation and related political decisions, we scrutinized the study and concluded that the main conclusions of Kubelka *et al*. (2018) are invalid (Bulla *et al*. 2019a). The authors then responded by reaffirming their conclusions (Kubelka *et al*. 2019b).

Here, we evaluate some of Kubelka *et al*.’s (2019b) responses, including their recent erratum (2020), and show that the main concerns about the original study still hold. Specifically, (1) we reaffirm that Kubelka *et al*.’s (2018) original findings are confounded by study site. Hence, their conclusions are over-confident because of pseudo-replication. (2) We reiterate that there is no statistical support for the assertion that predation rate has changed in a different way in the Arctic compared to other regions. The relevant test is an interaction between a measure of time (year or period) and a measure of geography (e.g., Arctic vs the rest of the world). The effect of such an interaction is weak, uncertain and statistically non-significant, which undermines Kubelka *et al*.’s (2018) key conclusion. (3) We further confirm that the suggested general increase in predation rates over time is at best a weak and uncertain trend. The most parsimonious hypothesis for the described results is that the temporal changes in predation rate are an artefact of temporal changes in methodology and data quality. Using only high-quality data, i.e. directly calculated predation rates, reveals no overall temporal trend in predation rate. Below we elaborate in detail on each of these points.

We conclude that (i) there is no evidence whatsoever that the pattern in the Arctic is different from that in the rest of the world and (ii) there is no solid evidence for an increase in predation rate over time. While we commend Kubelka *et al*. for compiling and exploring the data, we posit that the data underlying their study, and perhaps all currently available data, are not sufficient (or of sufficient quality) to test their main hypotheses. We call for standardized and consistent data collection protocols and experimental validation of current methods for estimating nesting success.

## (1) Multiple sampling and pseudo-replication

Each study site (and each year) has its own peculiarities that may affect the level of predation. For example, remote sites may have lower predator densities than a site close to a landfill. From our own experience in the Arctic, the presence of one or a few Arctic foxes (*Vulpes lagopus*) in a given year can lead to high predation rates on the nests of all studied shorebird species. These circumstances thus influence multiple datapoints from a study site. Such spatial non-independence in the data (the same fox eating the eggs of many species at a given site) is a form of ‘pseudo-replication’ and needs to be statistically controlled for; if not, the results may seem clearer than they actually are, leading to unsupported conclusions. Kubelka *et al*. (2019b) ascertain that their models (Kubelka *et al*. 2018) account for such multiple sampling per site, because “*spatial dependency is modeled as a random variance in which all points from the same location have the same (maximal) covariance. Our spatial term also accounts for possible covariance between sites that are close*.” We here show that this is not the case.

First, a simple mixed-effect model with only species as a random intercept, without any control for repeated sampling within sites, generates identical results to Kubelka *et al*.’s model that supposedly controlled for multiple sampling within sites (Table S1 in Bulla *et al*. 2019b). Kubelka *et al*.’s method would have controlled for a scenario where predation patterns are similar in nearby sites (spatial autocorrelation), but it does not control for multiple sampling per site (pseudo-replication). Indeed, Kubelka *et al*.’s (2018) way of controlling for spatial autocorrelation and multiple sampling within sites (the geographical distance matrix) explains none of the variance in random effects (Table S1A, C in Bulla *et al*. 2019b). Thus, this term has no influence on the model, which is equivalent to dropping it from the model.

Moreover, fitting the residuals from their original model (output in Table S2A in Kubelka *et al*. 2018) as the response variable and specifying site as a random intercept shows that site explains 57% of the variance. This effect differs between geographical belts, i.e. for models on subsets of the data (see Table S2B-F in Kubelka *et al*. 2018): site explains 0% of the variance for the South Temperate region, 93% for South Tropics, 70% for North Tropics, 55% for the North Temperate region, and 59% for the Arctic. The other original models (Table S6 in Kubelka *et al*. 2018) suffer from the same issue: site explains 55% of the variance in the residuals from the model on all data (Table S6a), 62% in the model on historic data (Table S6b), and 57% in the model on recent data (Table S6c). These results show unequivocally that Kubelka *et al*.’s (2018) method does not control for multiple sampling per site. The residuals of their models (presented in Tables S2 and S6) are not independent and show a non-random, site-dependent structure. In other words, Kubelka *et al*.’s models are heavily confounded by site and thus their results are statistically over-confident (confidence intervals around the estimate are too narrow) and pseudo-replicated.

This issue is not trivial, because – as we showed earlier – including ‘site’ as a random intercept reduces or wipes out the initially reported temporal trends (see Figure 1 in Bulla *et al*. 2019a). In response, Kubelka *et al*. (2019b) argued that having too few sites with multiple data points is problematic when controlling for pseudo-replication by including ‘site’ as a random intercept (for the exact quote see Note 1 below). While this generic argument might be valid in particular circumstances, it does not apply here. Of the 152 sites, 34 have more than one data point (Table R1) and of the six sites with more than five data points, three are from the Arctic. Also, 18 out of 49 Arctic sites have more than one data point (Figure R1). Importantly, our models robustly estimated the random term ‘site’, which explained 60-80% of the variance, as well as the residual term, which accounted for 20-30% of the variance (see the newly added 95% credible intervals for estimated random effects in Tables S2, S3, S5, and S6 in Bulla *et al*. 2019b). Thus, the argument of Kubelka *et al*. (2019b) is invalid, despite an erratum that supposedly fixed the issue (Note 2).

**Table R1.** Data distribution across sites.

| # of data points | 1 | 2 | 3 | 4 | 5 | 6 | 7 | 8 | 10 | 13 |
| --- | --- | --- | --- | --- | --- | --- | --- | --- | --- | --- |
| # of sites | 118 | 19 | 6 | 2 | 1 | 2 | 1 | 1 | 1 | 1 |

**Figure R1.**
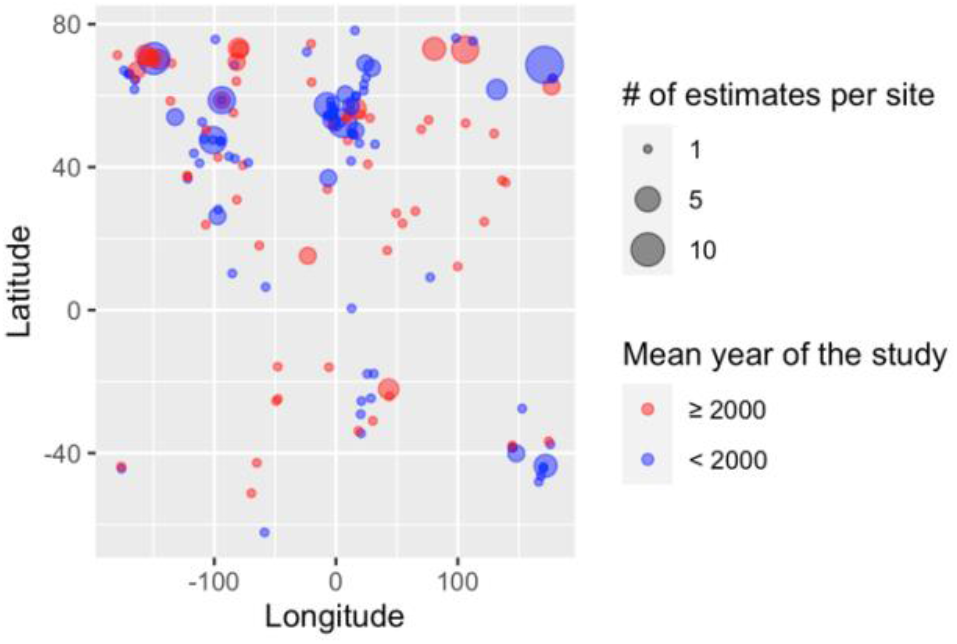
Geographical distribution of the study sites. Dot size indicates the number of predation rate estimates for a given site and dot color indicates whether the mean year of the study was before the year 2000 (blue) or not (red).

**Figure.**
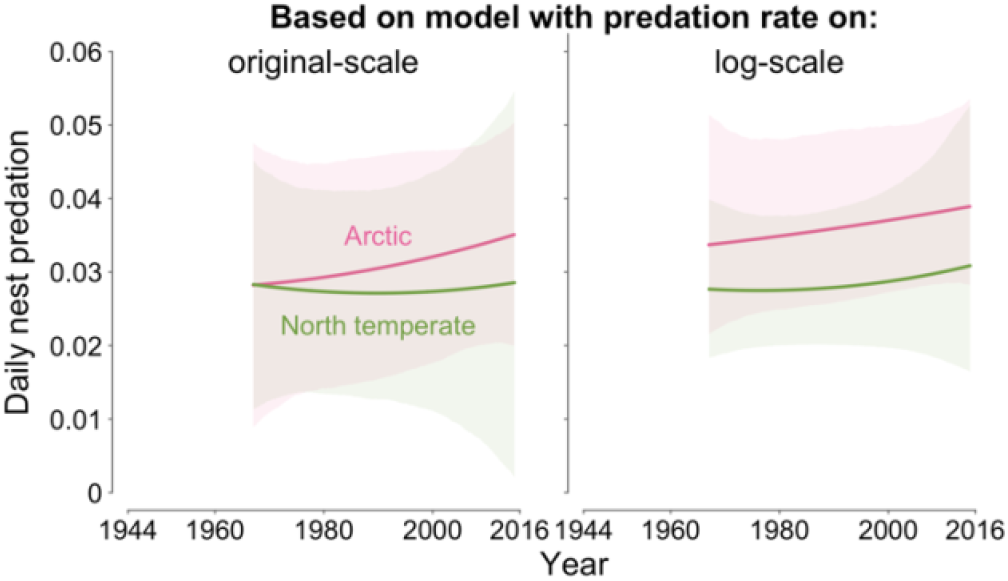
Original- and log-scale analyses show the absence of an interaction effect between year and geographical region. Comparison of predictions based on a model with daily predation rate on the original and log-transformed scale (for model specification see the Figure 1F legennd in Bulla *et al*. 2019a). Red indicates Arctic, green North temperate region. Note that given the wide 95%CIs around the predicted lines, the two methods give qualitatively identical results indicating no difference between the lines (regions) and hence no evidence for an interaction.

Kubelka *et al*. (2019b) further write *“When we removed groups represented by single observations* [and used only directly computed daily predation rates] *and reran the analysis [figure 1F in (2)], the temporal trend was indeed statistically significant using the approach of Bulla et al. (3)*.*”* We find this statement confusing because Kubelka *et al*. did not re-run our analysis (results of which are in Figure 1F in Bulla *et al*. 2019a). Instead, they ran an analysis that lacks a control for the latitude of the study site and for the number of nests used to calculate predation rate, and that does not include an interaction of time with latitude; doing so generates weak and statistically uncertain effects (see Note 3 for details).

Second, Kubelka *et al*. (2019b) claim that they controlled for pseudo-replication at the level of site, phylogeny and for the number of nests per population, but their figures are based on simple univariate general additive models (Figures 2A-D and 3 in Kubelka *et al*. 2018, or Figure 2C in Kubelka *et al*. 2019b) or linear models (Figure 1 in Kubelka *et al*. 2019b) that are neither controlled for site and phylogeny, nor for sample size, i.e. for the number of nests used to calculate a specific predation estimate. The same is true for reported means and standard errors in Figures 1, 2B and 2D in Kubelka *et al*. (2019b). The results shown in the figures are thus confounded by multiple sampling per site and by the number of nests per population (i.e. sample size), and hence do not provide adequate evidence for the claimed effects (see Figure R2 below).

**Figure.**
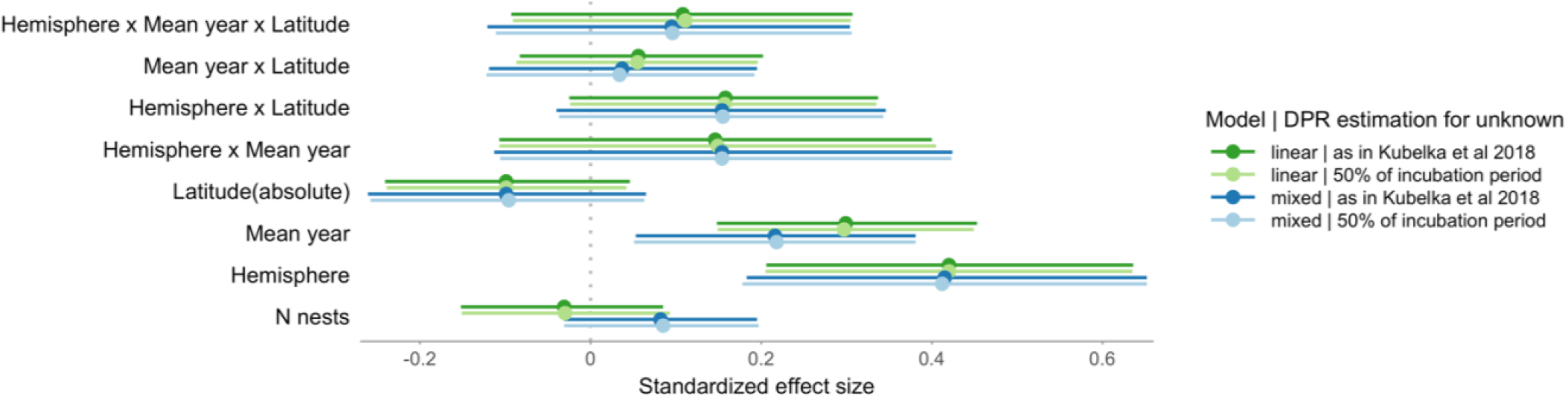
Sensitivity of the conclusions to the decision about how long nests in populations with unknown daily predation rates were followed. Dots with bars represent the estimates (medians) of the standardized effect sizes with the 95% credible intervals from a posterior distribution of 5,000 simulated values generated by the ‘sim’ function in R (Gelman and Su 2018). The response and all predictors were z-transformed (by subtracting the mean and dividing by the standard deviation). Before z-transformation the daily predation rates (after adding 0.01) and the number of nests (N nests) were log-transformed and ‘hemisphere’ was binarized (0 = South, 1 = North). N = 237 predation estimates representing 111 species and 152 sites. ‘As in Kubelka et al.’ (2018)’: data contain the same 50%, 60% and 90% transformation as in Kubelka et al. (2018); ‘50% of incubation period’: data are based on 50%-transformation for all data needing transformation; ‘linear’: simple linear models without control for multiple sampling within site; ‘mixed’: models controlled for site (random intercept). See Table below for further details.

**Figure.**
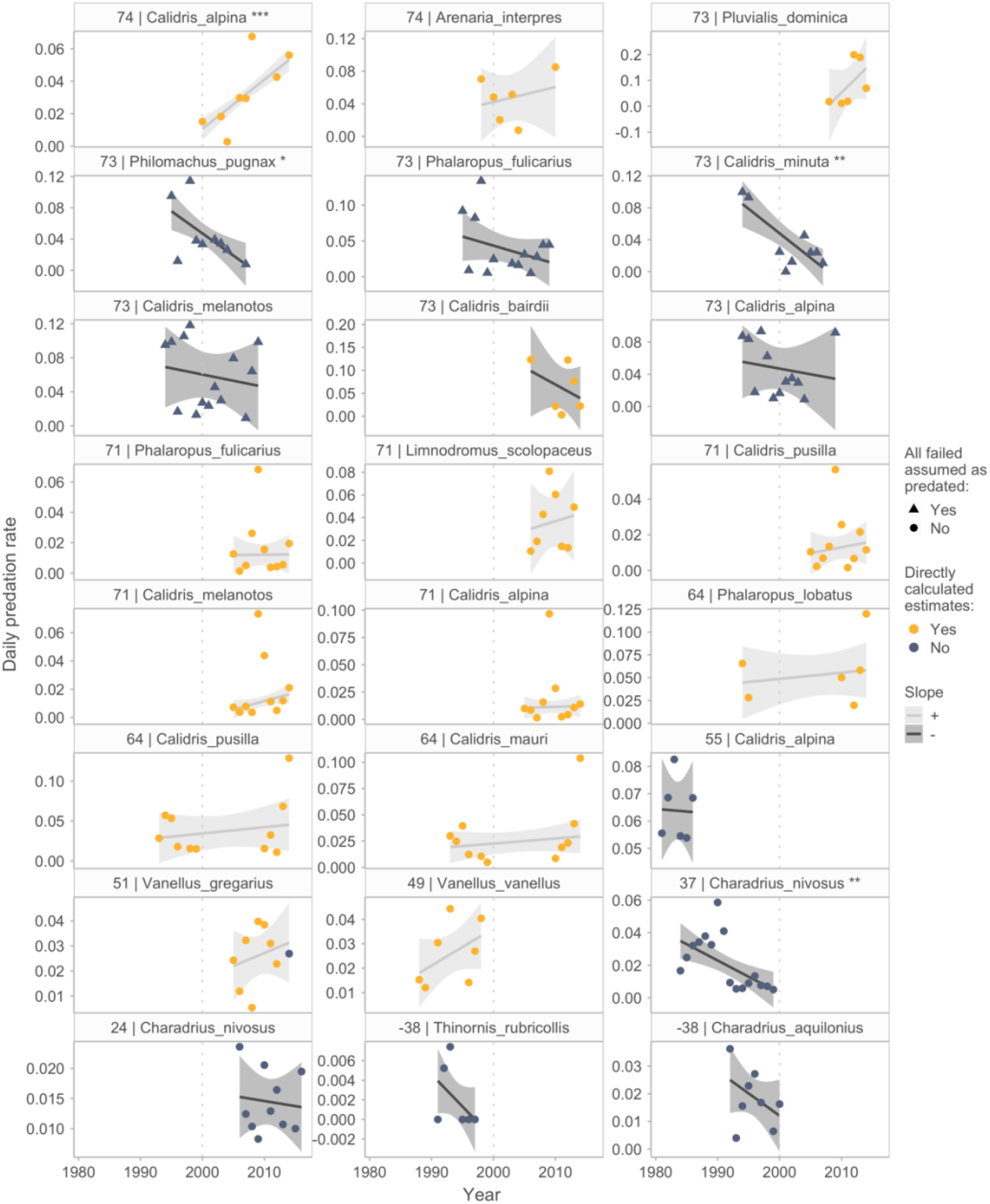
Between-population differences in temporal change in daily predation rates. Dots represent yearly estimates, their color indicates whether the estimate was directly calculated (yellow) or not (blue-grey), their shape whether the estimate is based only on predated nests (circles) or on all failed nests (triangles). Lines and 95%CIs are from robust regressions using M-estimators (Venables and Ripley 2002), not controlled for number of nests used to calculate each estimate, and generated by the ‘geom_smooth’ ggplo2 R-function. Note that robust regressions are less sensitive to outliers than linear regressions. The color of the lines and 95%CIs indicates whether the slope of the line is positive (light grey, N = 11) or negative (dark grey, N = 13; median slope: −0.01, mean: −0.04, range: −0.79 – 0.65; standardized effect size; 8 negative slopes after excluding populations with predation rate estimated based on all failed nests). Sites with positive and negative slopes occur in both periods (before and after 2000), but only four slopes were statistically significant (***p < 0.0001, ** p = 0.002 and 0.006, * p = 0.027); all other slopes had p > 0.25. Despite fox control in the 71°N site (Barrow, Alaska), predation rates varied between years with a weak increasing temporal trend. The dotted vertical line indicates year 2000 - Kubelka et al.’s (2018) divide between historic and recent data. The number in front of the species name indicates latitude of the study site. The panels are ordered from highest (top left) to lowest (bottom right) latitude and have a panel specific y-scale. Depicted are only years with more than 11 nests (following Kubelka et al.’s (2018) threshold) and only populations with more than five years of data.

Furthermore, Kubelka *et al*. suggest that their figures are supported by the provided model outputs. For example, the legend of Figure 2 in Kubelka *et al*. (2018) states *“model description* [for Figure 2A and B] *is available in table S2”*. However, this is not the case, because table S2 contains different models than those used to generate the figures (see scripts from Kubelka *et al*. 2018’s Dryad repository). As indicated above, Kubelka *et al*.’s (2018) scripts reveal that their figures are based on models that do not control for space, phylogeny or sample size. We also note that the figures in Kubelka *et al*. (2018) are based on models that do not test for the relationship of interest, namely the interaction between some measure of time and some measure of latitude, which leads to our next point.

## (2) Testing the key interaction effect: does the temporal change in predation vary across the globe?

The key conclusion of Kubelka *et al*. (2018) is that predation rates have increased over time more so in the Arctic than elsewhere. As we argued earlier (Bulla *et al*. 2019a), statistical evidence for this claim would be to show a significant interaction effect between time and region, with robust and biologically relevant effect sizes.

Kubelka *et al*. (2018) did test the interaction between year, absolute latitude and hemisphere, but do not report the weak, statistically non-significant results (p = 0.25 and p = 0.21, for daily and total predation rate respectively, Picture R1 below). Kubelka *et al*. (2018) did also test the interaction between year and absolute latitude (Table S6a in Kubelka *et al*. 2018) and report – but do not plot, nor discuss – the weak, statistically uncertain effects (p = 0.06 and 0.095; their Table S6A). Apart from not discussing these results in their original study (Kubelka *et al*. 2018), Kubelka *et al*. (2019b) also do not mention those in their response to our comment. Nevertheless, their results are consistent with our findings (Bulla *et al*. 2019a), although after controlling for study site (i.e. pseudo-replication) we found even weaker effects with no statistical support (Bulla *et al*. 2019b; p = 0.98 and p = 1 for the interaction between year and absolute latitude and p = 0.95 and p = 0.86 for the interaction between year, absolute latitude and hemisphere; Table R2 - A and B). Figures based on such models show unclear latitudinal or geographical differences (right panel in the Figure R2 below, Figure in Note 6 below, or Figure 1 in Bulla *et al*. 2019a).

The interaction between time and geographical region - untested, but claimed by Kubelka *et al*. (2018) - is statistically non-significant, both when controlling for multiple sampling per site or not (p = 0.65-0.96, Table R2 – C and D). Kubelka *et al*. (2019b) indicate that they have not tested this interaction between year and geographical region (or between year, absolute latitude, and hemisphere) and argue that it would be inappropriate to do so, because the response variable (daily predation rate) is log-transformed (see Notes 4 and 5). We consider this argument invalid, not only because Kubelka *et al*. (2018) did test for other interactions, e.g. one between year and absolute latitude discussed above (see Note 6 for details).

**Figure R2.**
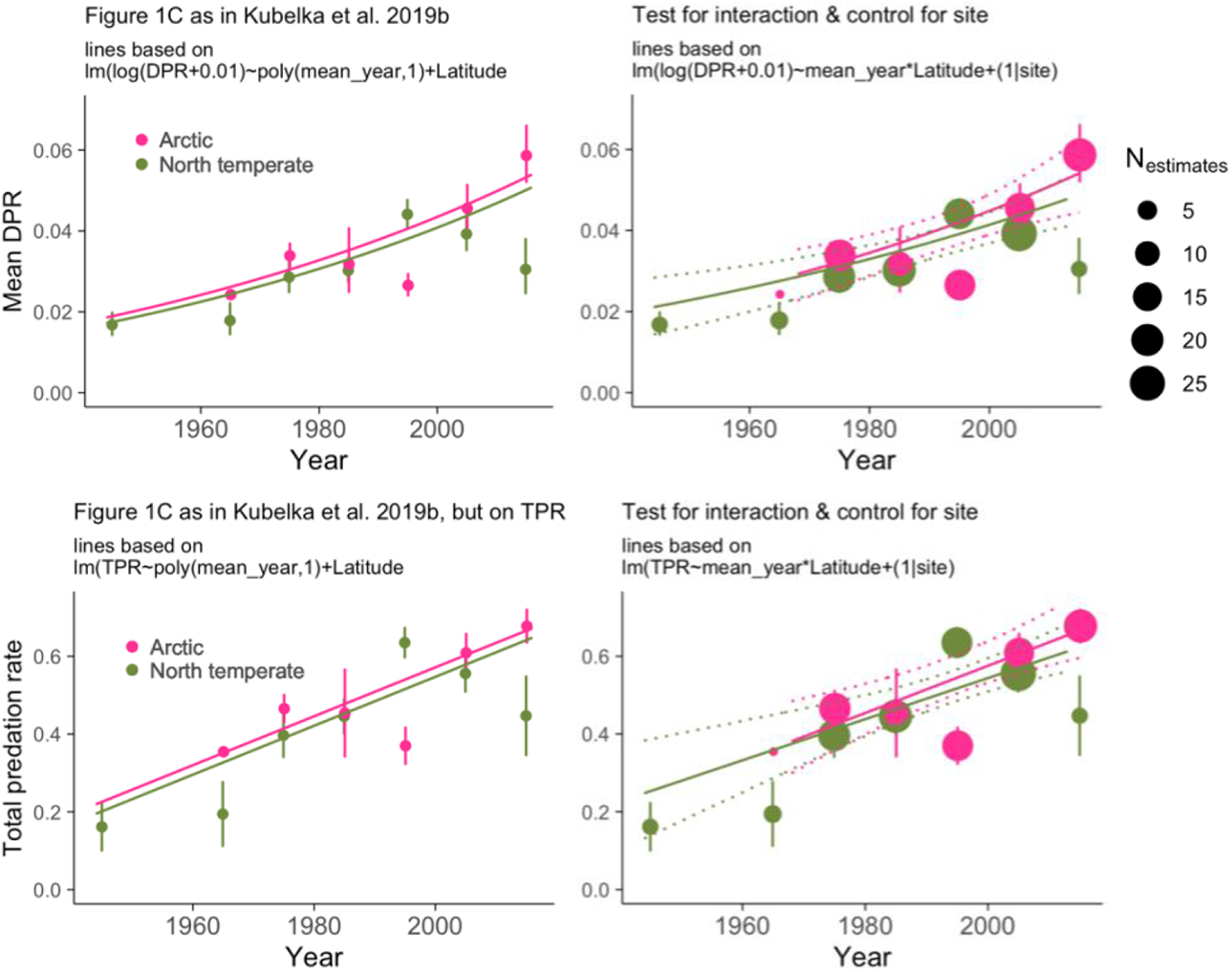
Scrutinizing Kubelka *et al*.’s (2019b) Figure 1. **Left panels** show a copy of Kubelka *et al*.’s (2019b) Figure 1C (top) and its reconstruction for total predation rate (bottom). Dots and bars represent decadal means ±SE and the lines show the predicted relationship for latitude at 69° (Arctic) and 50.7° (North temperate) from the linear model specified in the panels’ titles (see Kubelka *et al*.’s (2019a) script). Kubelka *et al*. (2019b) state in their Figure 1 legend “*both latitude and year are highly statistically significant…Note that although the model does not include an interaction term on the log scale, the lines diverge on the arithmetic scale, indicating geographically divergent outcomes*…” and elsewhere “*Differences between the Arctic and the nearest region (North temperate) are small on average, but recently daily nest predation has risen notably in the Arctic (Figure 1D* [based on Figure 1C]).” However, neither the decadal means, nor the predicted lines are controlled for sample size and pseudo-replication at site level, and hence cannot be used as evidence for a different effect in the Arctic. Kubelka *et al*. fitted latitude as a linear predictor (i.e. the effect of latitude reflects differences between Southern and Northern hemisphere as controlling this model for hemisphere renders the effect of latitude non-significant: p = 1). Essentially, the claimed geographically diverging outcomes are not reflected in the predicted lines for Arctic and North temperate (the two regions with the most data). Any apparent differences in such predicted lines (e.g. in their Figure 1A) are an artefact of data-transformation and not of a statistical interaction (which was not tested), i.e. the lines are parallel on a log-scale or when predation rate is fitted without transformation as is the case for total predation rate (bottom panel). **Right panels** show the reconstructed Figure 1C (left panels), but highlighting the differences in number of population estimates used to calculate each mean value (dot size) and showing predictions (solid lines) with 95% confidence intervals (dotted lines) based on the posterior distribution of 5,000 simulated values from a model (given in the panels’ titles) that tested for an interaction between mean year and latitude and controlled for multiple sampling per site. As in Figure 1C of Kubelka *et al*. (2019b), the predictions are for 69° latitude (Arctic) and for 50.7° (North temperate). Note that (a) the most recent difference in the means between Arctic and North temperate rests on a limited sample size for North temperate, and (b) the means just preceding the year 2000 show the opposite trend, and are based on more estimates; these issues are reflected in the weak predicted difference between Arctic and North temperate, with a non-significant interaction between latitude and year (p = 0.6, generated with ‘glht’ function from ‘multcomp’ package (Hothorn *et al*. 2008)). The results are even weaker if absolute latitude is used.

**Picture R1.**
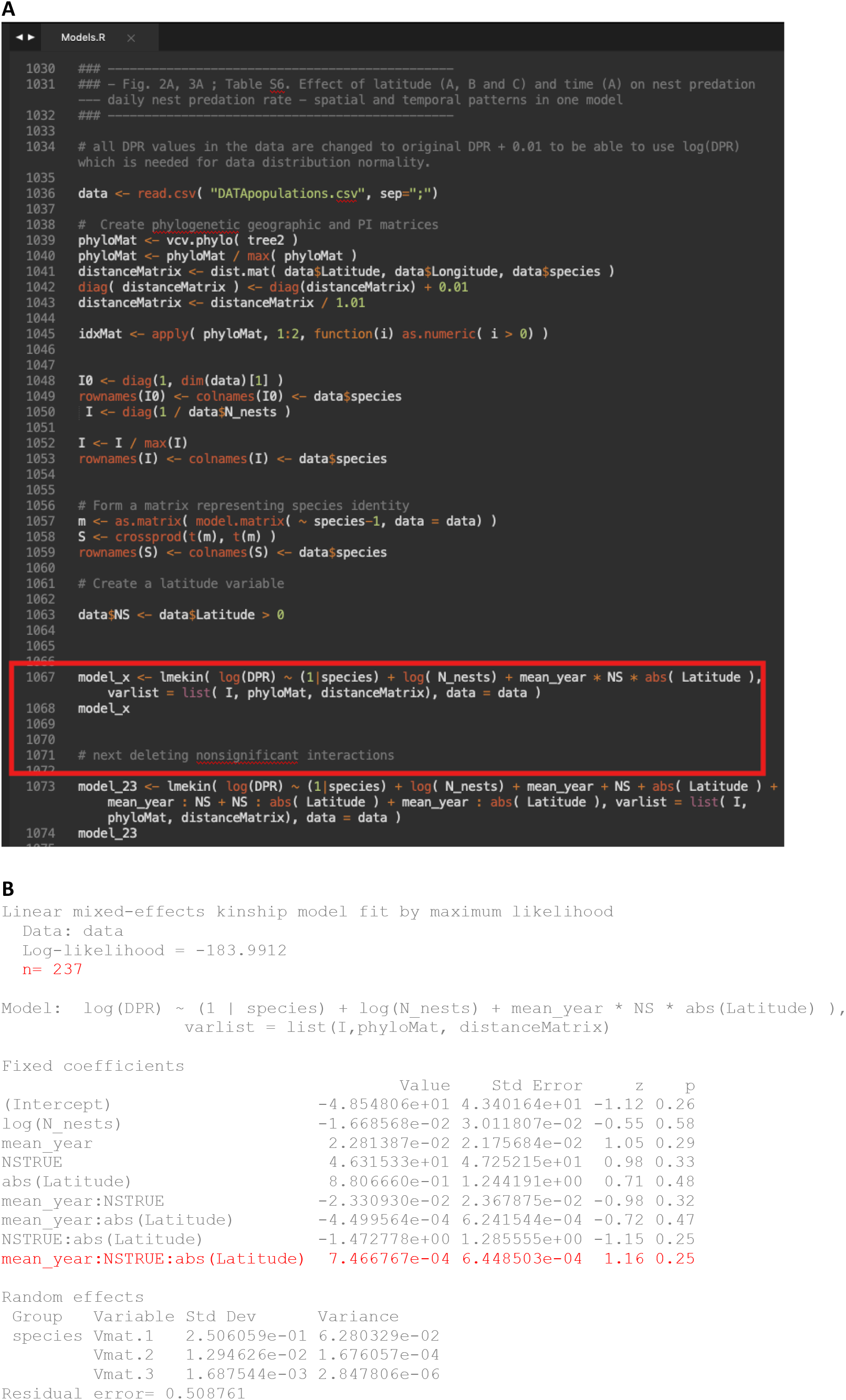
Kubelka *et al*.’s (2018) original script and the unreported results showing that the interaction of interest between year of the study, hemisphere and absolute latitude was statistically non-significant. **A**, Red rectangle highlights the unreported model of interest (see their Dryad script “Models.R”, line 1067 and 1288). Kubelka *et al*. (2018) also tested the interaction in the model on total predation rates (see line 1288 of their script). **B**, Output of the model highlighted in **A**. Note the weak, uncertain and statistically non-significant effect of the three-way interaction (in red). **A, B**, DPR = daily predation rate, N_nests = number of nests used to calculate a specific estimate, mean_year = mean year of the study, NS = hemisphere (and NSTRUE indicates Northern hemisphere) and Latitude = latitude of the study.

**Table R2.**
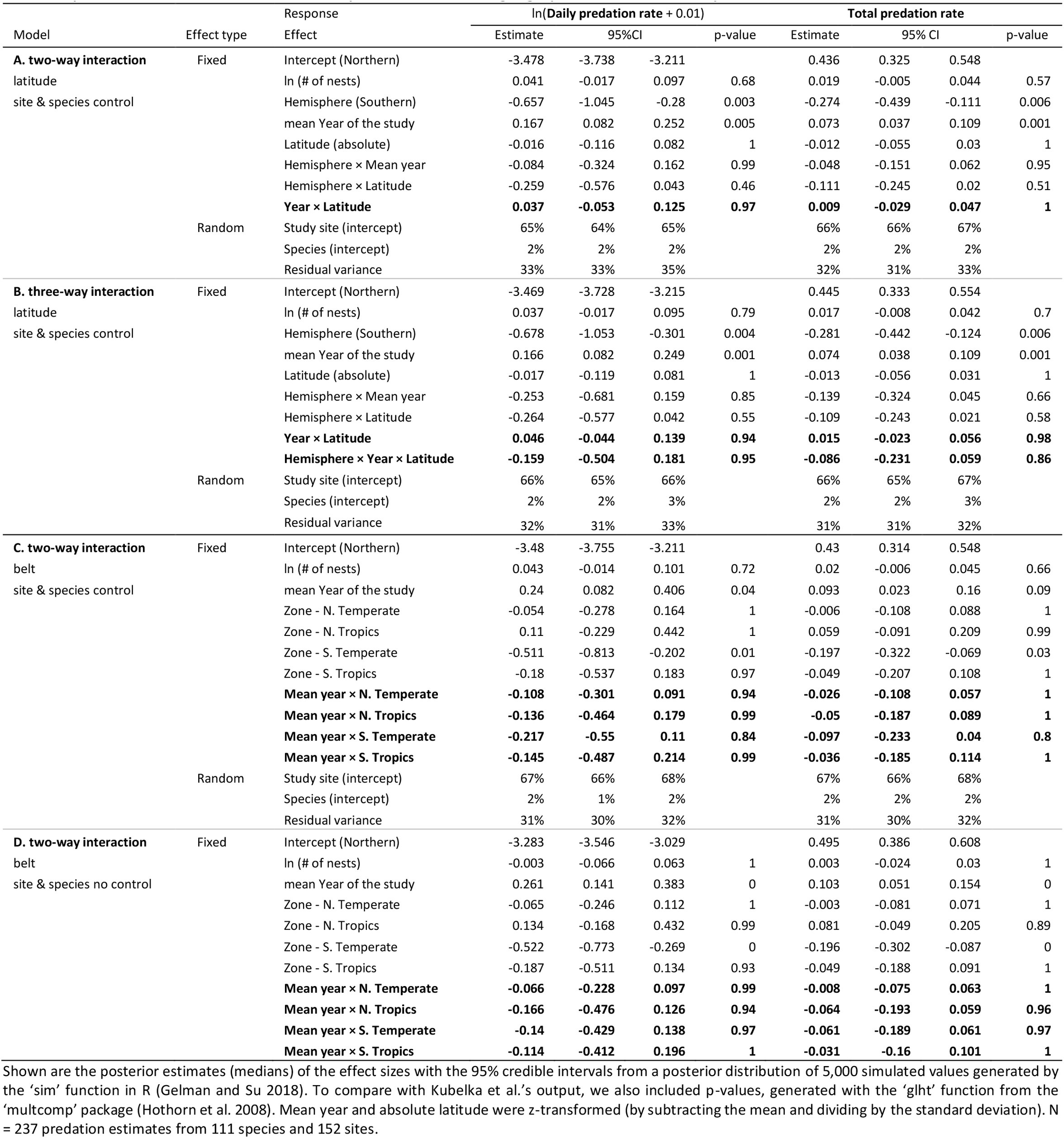
Predation rates in relation to mean year and latitude or geographical belt of the study.

## (3) Do predation rates increase over time?

Kubelka *et al*.’s (2018) crucial finding is that nest predation increased recently (globally, and especially in the Arctic). Here, we re-evaluate the evidence for this global increase over time and argue that it is unclear.

First, when we control for multiple sampling per site, the effect size for the general increase in predation rates over time is 30% lower than reported in Kubelka *et al*. (2018, Table S3). The effect size becomes essentially zero (0.005 [-0.006 - 0.016], p = 0.57) when we use only higher-quality, directly calculated predation rates (defined and discussed in Note 7), i.e. when we add the random effect ‘site’ to Kubelka *et al*.’s Table S3D model (for similar results from models containing also geographical region or latitude, see Tables S2, S3, S5, and S6 in Bulla *et al*. (2019b); for a further response to Kubelka *et al*.’s (2019b) arguments see Note 8).

Second, Kubelka *et al*. (2018; 2019b) used mean year of the study in their models. However, the underlying data cover periods of up to 44 years. Thirty estimates were based on 10 or more years and 18 of those contained data from both before and after the cut-off year (2000) used to define the early and late study period. For example, the data on the whimbrel (*Numenius phaeopus*) from Churchill stem from the years 1964-1967 (Jehl 1971), 1973-1974 (Skeel 1983) and 2011-2014 (Arctic-Shorebird-Demographic-Network 2016), but were used by Kubelka *et al*. (2018) for a predation estimate for the year 1990 (e.g. in their Figure 2A-D and Tables S2, S3 and S6a). It is unclear why Kubelka *et al*. (2018) did not use yearly estimates whenever they were available. Doing so reveals considerable between-population variability in temporal trends (Figure in Note 9) and leads to estimates of the ‘year’ effect that are practically zero (Table in Note 9).

Third, field and analytical methods have changed considerably over time, which makes the data difficult to compare. Essentially, methods for estimating predation have changed dramatically and especially so in the Arctic (Figure 1G in Bulla *et al*. 2019a; Figure S2 in Bulla *et al*. 2019b). Specifically, the number of studies that obtained more precise estimates of predation rate (defined and discussed in Note 7) increased over time, which may have biased the results in favor of the hypothesis, i.e. towards increasing predation rates over time. An apparent increase in predation rates may thus be an artefact of a change in methods. As we showed previously (Figure 1C and 1F in Bulla *et al*. 2019a), and here (e.g. Figure in Note 6), when only the data based on more precise estimates of predation rate are used, there is no evidence for a change over time.

We further hypothesized that the apparent temporal increase in nest predation could also be caused by an increase in research intensity over the years (Bridge *et al*. 2013; Dominoni *et al*. 2017; Andes *et al*. 2018) (reviewed in Warnock and Warnock 1993; Warnock and Takekawa 2003). Research intensity is a complex issue, because it not only involves nest searching, but also the use of nest and bird monitoring equipment. Yet, for example, information on nest-search intensity (also needed to estimate nest age at discovery) is not always clearly stated in the studies used in Kubelka *et al*. (2018) (see Note 7). Thus, research intensity may be hard to quantify. Kubelka *et al*. (2019b) deserve credit for addressing this issue: they “*scored the research intensity for the data used in (1)*.” Unfortunately, it remains unclear how this scoring was done, especially for the sources that lack the relevant information (see examples in the second paragraph of Note 7). It is also peculiar that when considering the different regions separately the research intensity score does not increase over time (Figure R3).

**Figure R3.**
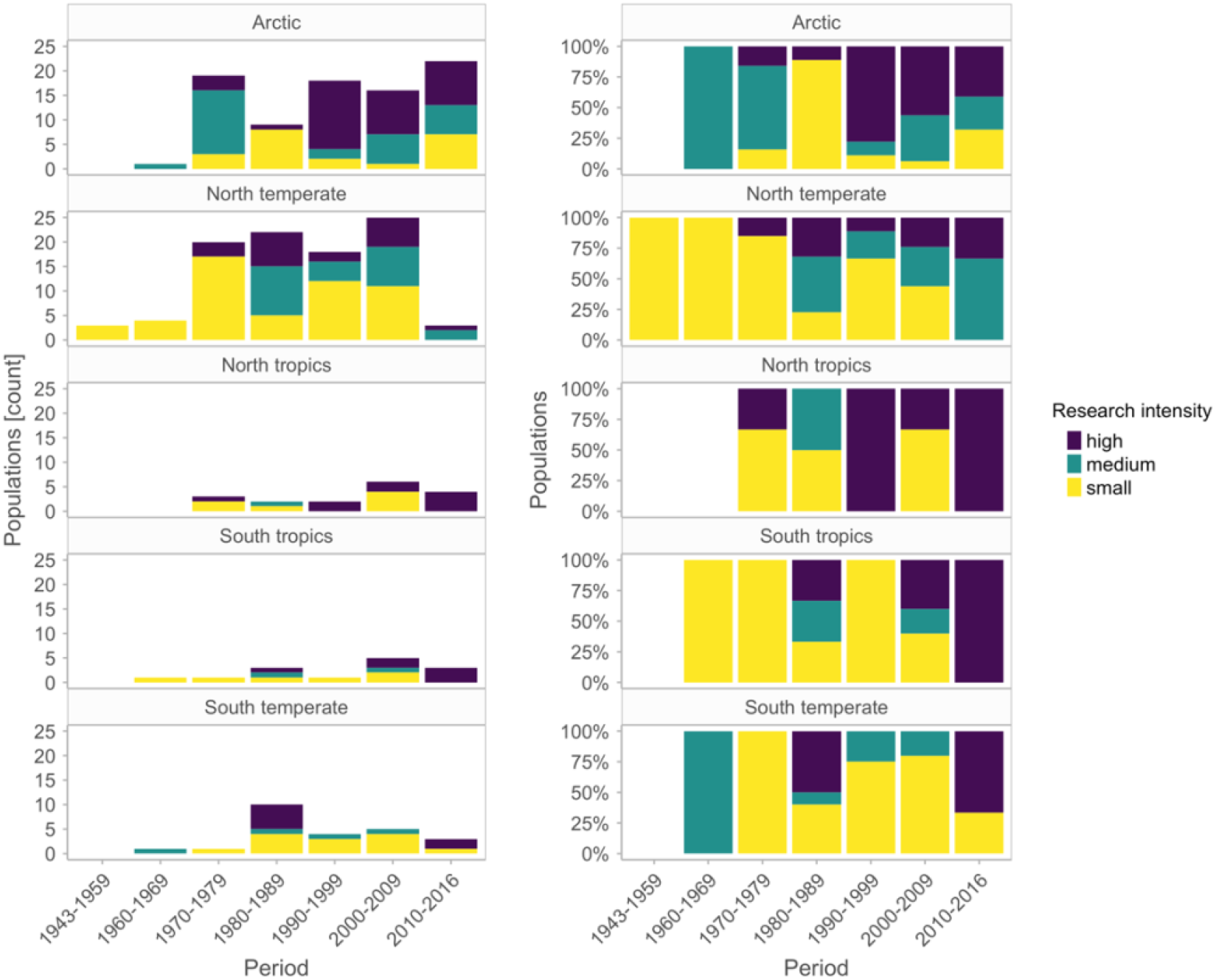
Distribution of Kubelka *et. al*’s (2019b) research intensity score across time and geographical region. Left panels represent counts, right panels corresponding percentages. Note the between-regions and between-periods stochasticity in research intensity.

## Conclusions and outlook

We show that the key problems with the original study outlined in Bulla *et al*. (2019a) remain valid despite the response by Kubelka *et al*. (2019b). The lack of control for repeated sampling per site and for changes in data quality over time, the use of mean year in long-term datasets to investigate temporal trends, and the difficulty to account for changes in the quality of predation estimates, as well as for an increased probability of observer-induced nest predation due to more invasive methods, all cast doubt on the conclusions of the original study. All these issues may bias the results in favor of the conclusion that there is a temporal increase in predation rates.

There is no clear evidence (1) for recent dramatic changes in predation rates in the Arctic nor (2) for a general increase in predation rates over time, as claimed by Kubelka *et al*. (2018; 2019b). At best, the evidence for a general increase is weak. We believe that the currently available data are unsuitable to answer the questions asked in the original study, because it is virtually impossible to control for changes in field and analytical methods, as well as in research intensity and invasiveness. This is not only true for Kubelka *et al*.’s study; other studies on nest predation suffer from similar issues (Remeš *et al*. 2012; Roodbergen *et al*. 2012).

To better understand variation and potential changes in nest predation rates, we advocate the use of standardized data collection protocols. Scientists should not only record the time during which each nest is observed, but also quantify disturbance. This can be done, for example, by continuously tracking the movements of all people doing fieldwork at a given study site with a handheld GPS and by clearly documenting the periods during which various equipment is installed on incubating birds or nests. If the focus of the study is on nest survival, field methods should preferably stay the same during the study period to allow unbiased assessment of temporal trends.

In the future, an important source of information about nest success may come from birds that have been equipped with tracking devices before the start of the breeding season. Nests can then be ‘found’ without visiting the area (Verhoeven *et al*. 2020) and survival estimates from such data can then be used and compared with those obtained via traditional methods.

We highlight that a focus on nest predation to explain declines in shorebird numbers may be misplaced.

## Code

All analyses are replicable with the open access code and material available from https://github.com/MartinBulla/Still_no_evidence/.

## Acknowledgements

We are grateful to Dov Lank, Fränzi Korner-Nievergelt, Brett Sandercock, Clemens Küpper, Krisztina Kupan, Wolfgang Forstmeier, Bruce Lyon, Bob Montgomerie, Jenny Gill, Anastasia Popovkina, Luke Eberhart-Hertel, Petr Keil, Emmi Schlicht, Martin Sládeček, and members of the Department of Behavioural Ecology & Evolutionary Genetics at the Max Planck Institute for Ornithology, as well as members of the Coastal Systems group at NIOZ for advice, comments and discussions. We are particularly grateful to the following signatories for help with revising this manuscript: Martijn van de Pol, Paul Smith, Eduardo SA Santos, Johannes Lang, Eunbi Kwon, Sarah E Jamieson, Nathan Senner, Marcelo Bertellotti, Nicolas Lecomte, Ricardo A. S. Cerboncini, Megan L Boldenow, Rose J Swift, Matthew Johnson, José A Alves, and Nicolas Meyer. We thank Vojtěch Kubelka, Miroslav Šálek, Pavel Tomkovich, Zsolt Vegvári, Robert P. Freckleton and Tamás Székely for feedback that helped improve this text.

## Notes

1. Kubelka *et al*. (2019b) argued that “*The large number of random group levels (111 species and 152 localities) used by Bulla et al*., *relative to the number of observations (237), as well as the large proportion of groups represented by just single observations (59 of 111 species and 114 of 152 localities), raise concerns about the robustness of their approach—specifically, the identifiability of the random and residual terms when the fixed effects vary at the level of localities*.
2. In response to our comments, Kubelka *et al*. noted that the variances for phylogeny and spatial terms were underestimated in their orginal model. They revised their R-scripts, and *Science* replaced the original Supplementary material with a new one containing new Tables. Kubelka *et al*. (2020) claim that the revised code “increased the variance attributable to space, accounting for multiple data points per field sites as well as distance among field sites”, but this is clearly not the case. The original models included only species as random intercept. The revised models include a random effect (*add code*) that has as many levels as there are rows in the dataset. This is puzzling and does not fix the problem of pseudo-replication. Using Kubelka *et al*.’s coxme model, but specifying site as a random intercept leads to the same conclusions as our mixed effect models: there is no support for the interaction of interest (using all data) or for the general increase in predation rates (using high quality data) - see here for details.
3. We specified the model as follows:

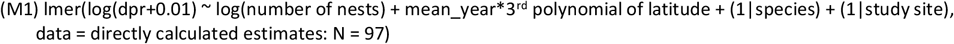 Kubelka *et al*. (2019b) fitted a model M2 that lacks (a) the interaction between time and latitude, (b) a polynomial fit for latitude (and thus also a control for latitude), and (c) a control for the number of nests used to calculate predation rate estimates (see line 278 of the script in Kubelka *et al*. 2019a; https://osf.io/46bt3/):

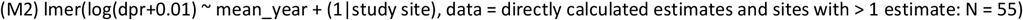 Indeed, the latter model (M2) generates a statistically significant year effect. However, the very same model (also included in Kubelka *et al*. 2019a, albeit unreported, see line 265 of their script) with all directly calculated estimates, i.e. estimates from data with known exposure (N = 97), as opposed to 55 in M2, yields a statistically non-significant year effect (model A in Table below). Importantly, the year effect is also weak and statistically non-significant when model M2 (on 55 data points) controls for the number of nests in a population and latitude of the study site (model B in Table below). Still such model does not test for the interaction of interest (as we did in M1), namely the interaction between time and latitude (see point 2 in the main text). Such interaction has little statistical support in the reduced dataset (N = 55) of Kubelka *et al*. (2019a) and the year effect is again weak and statistically non-significant (model C in Table below). Given that latitude in the reduced (N = 55) dataset spans only from 37° to 73°, fitting latitude as linear term might be more meaningful (model D in Table below), which also gives weak and statistically non-significant interaction and year effect.
4. Kubelka *et al*. (2019b) write *“Bulla et al. argue that we should have presented evidence of a statistically significant interaction between region/latitude and year/period on daily nest predation rate in order to demonstrate that the temporal trend of increasing predation rate varies spatially. This criticism ignores that the analyses both we and they reported are based on a logarithmically transformed response variable [log(x + 0*.*01): transformation applied based on model diagnostics]. The presence or absence of interactions based on transformed data is difficult to interpret. For example, log transformation means that on the arithmetic scale, the effects of model predictors are multiplicative, which implies interactive effects*.”
5. Whenever Kubelka *et al*. mention log-transformation, they mean ln-transformation. For consistency, we do the same. Also note that the log function in R means ln-transformation.
6. Kubelka *et al*. (2019b) write that *“The presence or absence of interactions based on* [log] *transformed data is difficult to interpret*.*”* This may well be true, but Kubelka *et al*. (2018) only log-transformed daily predation rates and analyzed total predation rates on the original scale (see Methods in their Supplementary material). Thus, testing and interpreting the interaction based on total predation rates is possible (see Figure R2 above). Analyses on non-transformed total predation rates generate similar results as analyses on log-transformed daily predation rates (figures in Kubelka *at al*. (2018) and Table 1 and S1-6 in Bulla *et al*. 2019a, b). Importantly, as we discuss here (see Picture R1 above or Table S6 in Kubelka *at al*. 2018), Kubelka *et al*. (2018) have tested interactions, despite log-transformation of daily predation rates. Furthermore, Kubelka *et al*. (2019b) argue that the “*log(x + 0*.*01) transformation [was] applied based on model diagnostics”*. However, analyses of daily predation rates on the original and the log-transformed scale generate similar results (see Figure below). Also, note that the graphical representations of Kubelka *et al*.’s (2018) results (e.g. their Figure 2A-D and 3) are based on simple univariate general additive models, in which daily predation rate was used on the original (non-transformed) scale. In addition, Kubelka *et al*. (2019b) state that “*log-transformation means that on the arithmetic scale, the effects of model predictors are multiplicative, which implies interactive effects*.” When the response variable is log-transformed, the effects of model predictors are indeed multiplicative on an arithmetic scale. However, this does not imply a statistical interaction, i.e. a statistical difference between the slopes. Kubelka *et al*. (2019b) argue their case by stating “*The simple model reported in Figure 1A captures this* [geographic difference between regions in] *increase* [through time], *despite the lack of an interaction term, because of the data transformation*.” However, the difference between the estimated lines for the different regions is not clearly visible in their Figure 1A or 1C (Kubelka *et al*. 2019b) – see reconstructed Figure 1C in our Figure R2 above. Moreover, the lines in Kubelka *et al*.’s (2019b) Figure 1A and 1C are generated from a model that does not control for sample size and multiple sampling per site (see Picture below) and such lines are parallel on the log-scale, i.e. the scale on which the data were analyzed (see Figure R2 above). Further note that the initial claim was about the Arctic being different from other regions, not whether the N hemisphere differs from the S hemisphere (the latter with limited sample size), although the interaction between mean year and hemisphere is clearly non-significant (p = 0.9), regardless of whether we control for study site or not.

**Picture.**
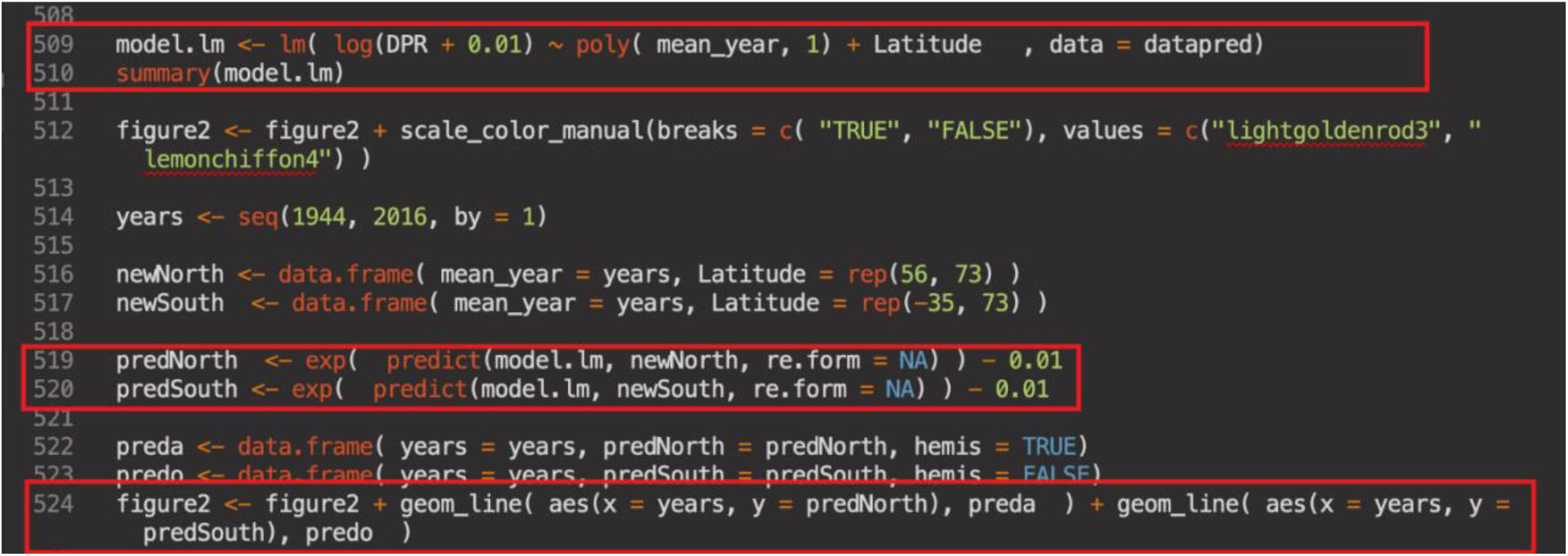
Kubelka *et al*.’s (2019a) script shows that Figure 1A in Kubelka *et al*. (2019b) is based on a model that does not control for multiple sampling per site, nor for the number of nests used to generate specific estimates. The same model is used and hence the same issues hold for Figure 1C (see lines 407-445 of their script and Figure R2 above).
7. Kubelka *et al*. (2018) calculated daily nest predation rates as the number of depredated nests divided by ‘exposure’ [the total time (in days) all nests were observed]. For the 59% of the 237 populations where they lacked information on exposure, they estimated exposure based on the description of nest search intensity in the respective studies. The key question is when nests were found. Kubelka et al. decided that in 114 populations, nests were found such that 60% of the nesting period (egg laying and incubation combined) had been ‘observed’ (nests searched once or twice a week). For 14 populations they used 90% (nests searched daily or found just after laying), and for 11 populations they used 50% (assuming nests found midway during the nesting period). However, it is unclear whether such assumptions are valid. Importantly, as indicated in our comment (Bulla et al. 2019a), Kubelka *et al*.*’*s (2018) sources (most of those available in our repository) do not always contain clear information about nest searching and nest checking intensity, or about when within the incubation period nests were found – information essential for determining the % of the incubation period for which nests in populations with unknown (as well as known) daily-predation rates were followed (e.g. Jehl 1971; Brosset 1979 in French; Kondratjew 1982 in Russian; Reid and Montgomerie 1985; Moitoret *et al*. 1996; refs #137, #207, #133 and its translation, #217, #58 in Kubelka *et al*. 2018). In other words, the decision about the ‘% of the incubation period followed’ is often subjective. Similar issues hold for research intensity (e.g. Reid and Montgomerie 1985). Moreover, the sources sometimes lack clear information about the number of predated nests (e.g. Davis 1994; refs #165). Note that it remains unclear how sensitive Kubelka *et al*.’s (2018) conclusions are to a decision about whether 50%, 60% or 90% of the incubation period is assigned to a population. Assigning 50% (the default value) to all data without directly calculated daily predation estimates (i.e. also those where Kubelka *et al*. (2018) assigned 60% and 90%) has little influence on the estimates (see Figure below). Thus, the methodological bias does not seem to arise from the decision about the %-value used to calculate daily predation rates, but rather from the general need to transform part of the data. This need for transformation changes over time (Figure 1G in Bulla et al. 2019a) in a similar manner as the estimated predation rate.

**Table.** Scrutinizing Kubelka *et al*.’s (2019b) replication of our analyses.

| Model | Effect type | Response<br>Effect | ln(Daily predation rate + 0.01) |  |  | p-value |
| --- | --- | --- | --- | --- | --- | --- |
|  |  |  | Estimate | 95%CI |  |  |
| **A.**<br>N = 97 observations<br>from 61 sites | Fixed | Intercept (Northern) | -13.306 | -34.797 | 7.653 |  |
|  |  | **mean Year of the study** | **0.005** | **-0.005** | **0.016** | **0.34** |
|  | Random | Study site (intercept) | 77% | 78% | 76% |  |
|  |  | Residual variance | 23% | 22% | 24% |  |
| **B.**<br>N = 55 observations<br>from 19 sites | Fixed | Intercept (Northern) | -31.565 | -68.936 | 6.496 |  |
|  |  | ln (# of nests) | 0.088 | 0.012 | 0.17 | 0.08 |
|  |  | **mean Year of the study** | **0.013** | **-0.006** | **0.033** | **0.4** |
|  |  | Latitude (absolute) | 0.019 | -0.012 | 0.049 | 0.52 |
|  | Random | Study site (intercept) | 77% | 78% | 76% |  |
|  |  | Residual variance | 23% | 22% | 24% |  |
| **C.**<br>N = 55 observations<br>from 19 sites | Fixed | Hemisphere × Latitude | -3.068 | -4.037 | -2.051 | 0 |
|  |  | ln (# of nests) | 0.096 | 0.015 | 0.178 | 0.06 |
|  |  | **mean Year of the study** | **0.976** | **-0.007** | **1.959** | **0.14** |
|  |  | Latitude (1<sup>st</sup> polynomial) | -10.585 | -26.125 | 5.271 | 0.46 |
|  |  | Latitude (2<sup>nd</sup> polynomial) | 12.272 | -5.949 | 30.926 | 0.46 |
|  |  | Latitude (3<sup>rd</sup> polynomial) | -2.955 | -7.596 | 1.55 | 0.47 |
|  |  | **Year × Latitude (1<sup>st</sup> polynomial)** | **-13.144** | **-34.01** | **7.619** | **0.5** |
|  |  | **Year × Latitude (2<sup>nd</sup> polynomial)** | **17.909** | **-6.281** | **42.294** | **0.38** |
|  |  | **Year × Latitude (3<sup>rd</sup> polynomial)** | **-3.988** | **-12.42** | **4.089** | **0.7** |
|  | Random | Study site (intercept) | 76% | 74% | 77% |  |
|  |  | Residual variance | 24% | 23% | 26% |  |
| **D.**<br>N = 55 observations<br>from 19 sites | Fixed | Intercept (Northern) | -3.603 | -4.054 | -3.168 |  |
|  |  | ln (# of nests) | 0.087 | 0.006 | 0.169 | 0.13 |
|  |  | **mean Year of the study** | **0.169** | **-0.092** | **0.434** | **0.61** |
|  |  | Latitude | 0.194 | -0.102 | 0.496 | 0.62 |
|  |  | **Year × Latitude** | **0.077** | **-0.273** | **0.424** | **0.99** |
|  | Random | Study site (intercept) | 78% | 79% | 78% |  |
|  |  | Residual variance | 22% | 21% | 22% |  |

Output of model M2 after using all directly calculated predation rate estimates (A), after using only populations with multiple observations, while controlling for the number of nests per predation estimate and for latitude of the study site (B), and after including the interaction of interest as in our original M1 model (C), which Kubelka *et al.* (2019b) attempted to replicate, as well as with linear latitude because latitude in the reduced dataset ranges only between 37° to 73° (D). Highlighted (in bold) is the effect of year (i.e., overall temporal change in predation rate) and of the interaction of interest, both of which are consistently statistically non-significant. Shown are the posterior estimates (medians) of the effect sizes with the 95% credible intervals from a posterior distribution of 5,000 simulated values generated by the ‘sim’ function in R (Gelman and Su 2018). To compare with Kubelka *et al.*’s output, we also included p-values, generated with the ‘glht’ function from the ‘multcomp’ package (Hothorn et
al. 2008). In the models with interaction (C, D), mean year and absolute latitude were z-transformed (by subtracting the mean and dividing by the standard deviation).

**Table.**
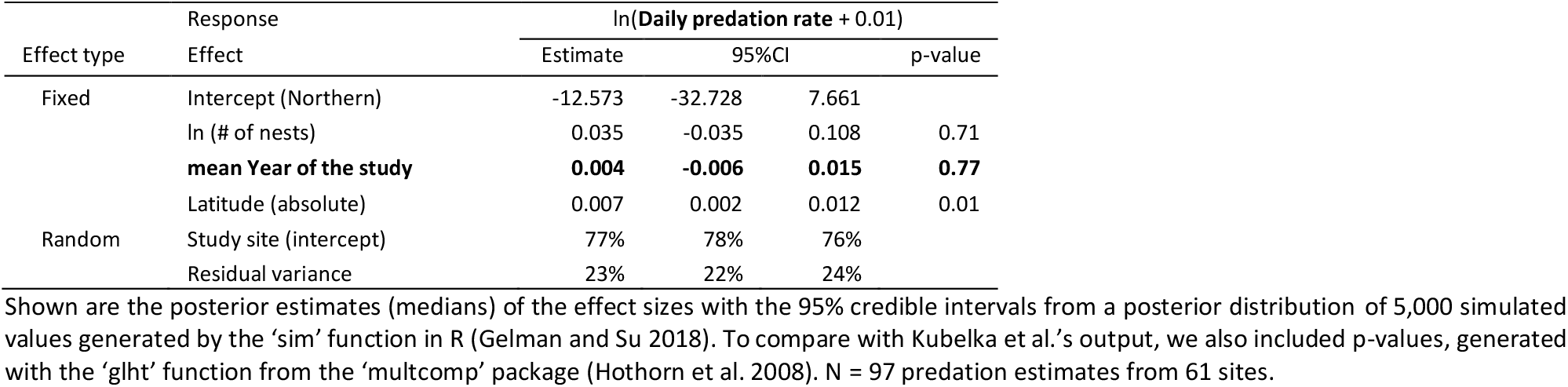
Output from model M4 that controls for multiple sampling within study site (note the non-significant year effect in bold)

**Table.** Predation rates in relation to year of the study using yearly estimates and populations with >5 years of data (as in the above Figure)
8. Kubelka *et al*. (2019b) write *“Contrary to Bulla et al*., *we find that the global temporal increase in nest predation is significant when daily nest predation was either calculated directly or converted from “apparent predation” [table S3 in (1); Fig. 2C] (14)*”. The models in Table S3 of Kubelka *et al*. (2018) are not controlled for study site (i.e. multiple data points per site) and doing so reduces the global temporal increase by 30% and for directly calculated daily nest predation renders the effect non-significant (0.005 [-0.006 - 0.016], p = 0.57)). The discussed Figure 2C (Kubelka et al. 2019b) is generated from univariate general additive models with year as the sole predictor, i.e. the results were not controlled for the number of nests per population or for multiple data points per site or species and latitude of the study. Moreover, Kubelka *et al*.’s (2019b) script (Kubelka *et al*. 2019a) – in addition to model M2 (Note 3) and the model shown in Picture R1A (in the main text) – provides the following model that could further support the above Kubelka *et al*. (2019b) *statement:*

| Model | Effect type | Effect | Estimate | 95%CI |  | p-value |
| --- | --- | --- | --- | --- | --- | --- |
| **A. All estimates** |  |  |  |  |  |  |
| N = 221 estimates<br>36 years<br>12 sites & 17 species | Fixed | Intercept | -3.366 | -3.537 | -3.19 |  |
|  |  | log (# of nests) | -0.081 | -0.179 | 0.02 | 0.39 |
|  |  | Latitude | 0.215 | 0.074 | 0.359 | 0.02 |
|  |  | **Year** | **-0.047** | **-0.215** | **0.121** | **0.97** |
|  | Random | Study site (intercept) | 0% | 0% | 0% |  |
|  |  | Year (slope) | 10% | 5% | 15% |  |
|  |  | Species (intercept) | 11% | 6% | 17% |  |
|  |  | Residual variance | 79% | 68% | 89% |  |
| **B. Estimates based on predation only** |  |  |  |  |  |  |
| N = 164 estimates<br>35 years<br>11 sites & 15 species | Fixed | Intercept | -3.417 | -3.632 | -3.202 |  |
|  |  | log (# of nests) | -0.094 | -0.208 | 0.022 | 0.37 |
|  |  | Latitude | 0.225 | 0.037 | 0.413 | 0.06 |
|  |  | **Year** | **0.009** | **-0.183** | **0.209** | **1** |
|  | Random | Study site (intercept) | 0% | 0% | 0% |  |
|  |  | Year (slope) | 18% | 10% | 26% |  |
|  |  | Species (intercept) | 9% | 5% | 13% |  |
|  |  | Residual variance | 73% | 61% | 86% |  |

Shown are the posterior estimates (medians) of the effect sizes with the 95% credible intervals from a posterior distribution of 5,000 simulated values generated by the 'sim' function in R (Gelman and Su 2018). To compare with Kubelka *et al.*'s output, we also included p-values, generated with the 'glht' function from the 'multcomp' package (Hothorn *et al.* 2008). Daily predation rates was log-transformed (after adding 0.01). To improve model convergence, the predictors were z-transformed (mean centered and divided by the standard deviation). We allowed for different temporal slopes within study site (i.e. accounted for the non-independence of data points that pseudo-replicate year-slope information). Models without such random slopes generated similar results. **A**, Data contain all estimates from the above Figure, i.e. population with >5 years of yearly estimates. **B**, Data as in **A**, but excluding populations with predation rates estimated based on all failed nest (see shapes in the Figure above).

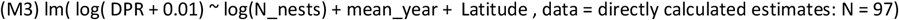 However, this model (M3) does not control for multiple sampling per site and adding this control (M4) renders the temporal trend statistically non-significant (see Table below or Figure F in Bulla et al. 2019a).

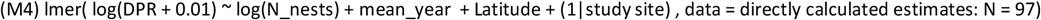
9. Bulla *et al*. (2019a) re-extracted most of Kubelka *et al*.’s data and where possible extracted yearly predation estimates that were based on more than 11 nests. The predation rates varied greatly within populations and between-population temporal trends were inconsistent (see Figure below). We found little support for a strong general increase in predation rates over time (see Table below). Note that Kubelka *et al*. (2019b) question the quality of the re-extracted dataset. Although it is possible that the re-extracted dataset contains some errors, these are unlikely systematic. Any discrepancies due to subjective decision about conversion of apparent predation rates to daily predation rates shall have negligible effects on the below reported results because use of different conversion coefficients has minor influence on the findings (see Figure in Note 7) and because we investigated within population changes; within each population, predation rate estimation methods are uniform (see Figure below). Importantly, Kubelka *et al*.’s data are not mistake free. For example, Kubelka *et al*. (2018) used only part of the freely available data (e.g. from ASDN database or from Soloviev *et al*.’s study in Southeastern Taimyr).

---

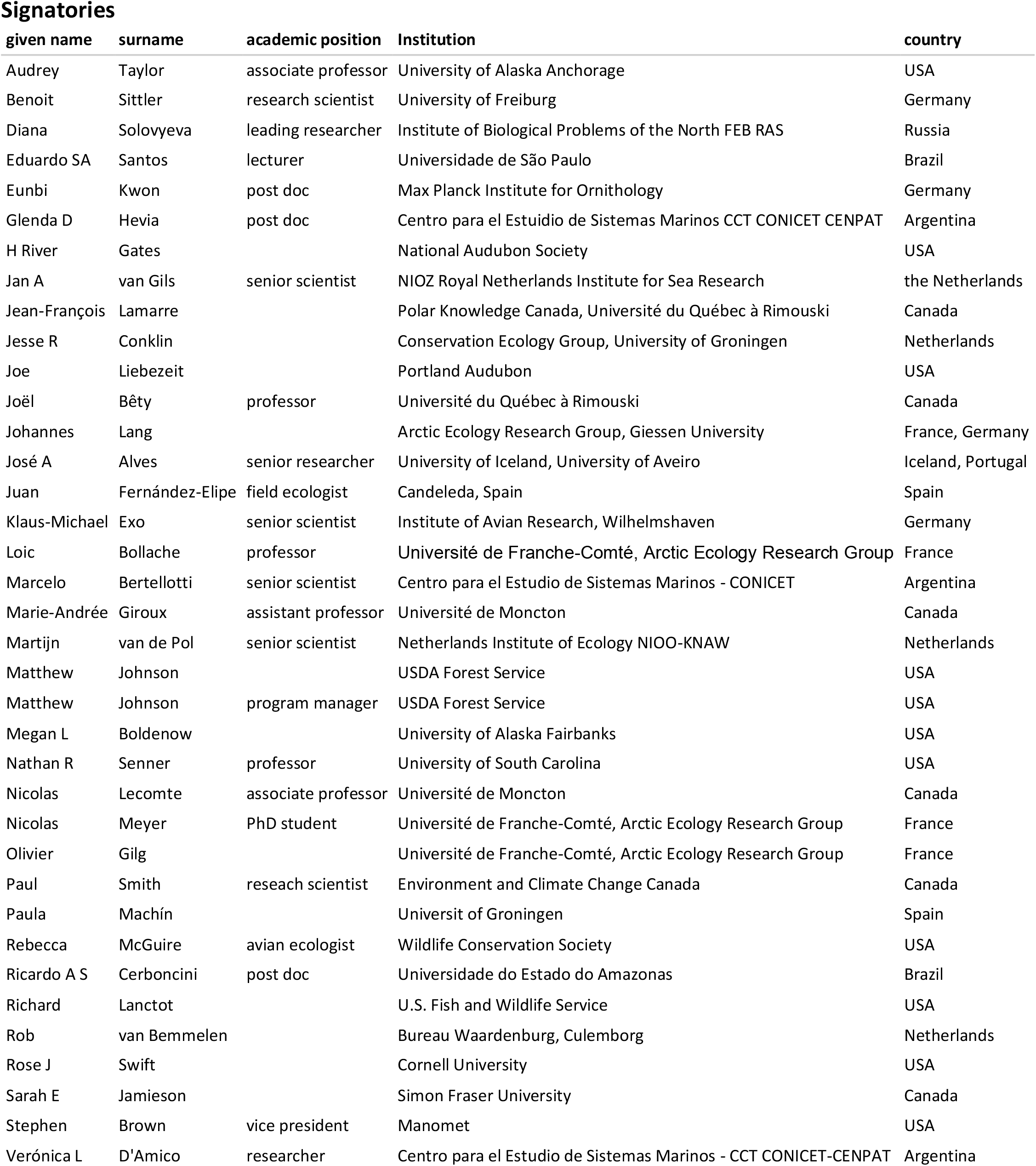

